# Long-term taxonomic and functional divergence from donor bacterial strains following fecal microbiota transplantation in immunocompromised patients

**DOI:** 10.1101/109645

**Authors:** Eli L. Moss, Shannon B. Falconer, Ekaterina Tkachenko, Mingjie Wang, Hannah Systrom, Jasmin Mahabamunuge, David A. Relman, Elizabeth L. Hohmann, Ami S. Bhatt

**Affiliations:** Departments of Genetics, and Medicine, Stanford University, Stanford, California, USA.; Departments of Microbiology and Immunology, and Medicine, Stanford University, Stanford, California, USA.; Division of Infectious Diseases, Massachusetts General Hospital; Harvard Medical School, Boston, Massachusetts, USA.; Veterans Affairs Palo Alto Health Care System, Palo Alto, California, USA.

## Abstract

Immunocompromised individuals are at high risk of developing *Clostridium difficile-*associated disease (CDAD). Fecal microbiota transplantation (FMT) is a highly effective therapy for refractory or recurrent CDAD and, despite safety concerns, has recently been offered to immunocompromised patients. We investigated the genomics of bacterial composition following FMT in immunocompromised patients over a 1-year period. Metagenomic, strain and gene-level bacterial dynamics were characterized in two CDAD-affected hematopoietic stem cell (HCT) recipients following FMT. We found alterations in gene content, including loss of virulence and antibiotic resistance genes. These alterations were accompanied by long-term bacterial divergence at the species and strain levels. Our findings suggest limited durability of the specific bacterial consortium introduced with FMT and indicate that alterations of the functional potential of the microbiome are more complex than can be inferred by taxonomic information alone. Our observation that FMT alone cannot induce long-term donor-like alterations of the microbiota of HCT recipients suggests that FMT cannot indefinitely supersede environmental and/or host factors in shaping bacterial composition.

## Introduction

Fecal microbiota transplantation (FMT) is a remarkably safe and effective therapy for resolving recurrent *Clostridium difficile*-associated disease (CDAD) in immunocompetent individuals (van Nood et al. 2013), and it is increasingly being used to treat CDAD in immunocompromised individuals (Kelly et al. 2014). Among immunocompromised populations at greatest risk of developing CDAD are hematopoietic stem cell transplantation (HCT) recipients, who experience attack rates as high as 25%, and who are up to nine times more likely to suffer from CDAD than immunocompetent individuals (Bruminhent et al. 2014; Chopra et al. 2011; de Castro et al. 2015). A small number of recent studies have reported that FMT can safely resolve CDAD in post-HCT recipients (Kelly et al. 2014; Mittal et al. 2015; Neemann et al. 2012; Aroniadis et al. 2016; Webb et al. 2016). However, persisting concerns regarding the safety of administering FMT therapy to immunocompromised patients continue to limit use of the therapy in post-HCT individuals. Better understanding the effects of FMT through investigations of post-treatment gut bacterial community structure in immunocompromised patients is needed.

Several studies leveraging 16S rRNA gene sequencing have begun to reveal the taxonomic underpinnings of FMT therapy by informing on genus- and species-level bacterial dynamics. These studies have demonstrated that FMT induces taxonomic compositional changes in the recipient gut microbiota, resulting in a more donor-like state (Seekatz et al. 2014; Shahinas et al. 2012; Hamilton et al. 2013; Broecker et al. 2016; Jalanka et al. 2016). These analyses have provided a foundation for metagenomic investigation of FMT: there are good reasons for considering whole genomes and strain-level diversity, since bacterial strains with identical 16S rRNA sequences can exhibit wide variability in terms of phenotype and pathogenicity resulting from nucleotide level DNA sequence differences or differing gene content. For example, commensal *Escherichia coli* are abundant facultative anaerobes in the human gut (Tenaillon et al. 2010), yet enterohemorrhagic *Escherichia coli* are important pathogens. A recent study of patients undergoing FMT for treatment of metabolic syndrome was the first to report strain-level dynamics after FMT. In this study of 10 patients, the investigators found that strains of donor bacteria are capable of both replacing and coexisting with same-species recipient strains at three months post-FMT (Li et al. 2016). These findings are intriguing, but questions remain regarding the generalizability of this finding to patients undergoing FMT for treatment of CDAD where, unlike the metabolic syndrome cohort, patients receive antibiotics prior to FMT and have an underlying dysbiosis. In the setting of prior antibiotic therapy, it is presumed that the total amount and diversity of microorganisms present in the FMT recipient is lower, and thus the gut may represent a relatively barren ecological niche for repopulation with donor-derived microbes. Additionally, no studies have yet reported on strain level engraftment at time points greater than three months after FMT.

To date, the collective understanding of FMT impact on the taxonomic composition of the gut microbiota has been defined by studies of immunocompetent patients. However, as immunocompromised patients are at greatest risk of developing CDAD, there is much to gain from an investigation of FMT in this patient population, in which the impaired host immune system may have diminished influence on the gut microbiota community. Further, although the recent studies relating the success of FMT in immunocompromised individuals have begun to assuage concerns of safety (Kelly et al. 2014; Mittal et al. 2015; Neemann et al. 2012; Aroniadis et al. 2016; Webb et al. 2016), those concerns are not unfounded (Quera et al. 2014). Characterizing the taxonomic and functional capacity of the immunocompromised patient microbiota post-FMT carries potential for both improving clinical outcomes of CDAD in patients lacking a robust immune system and advancing our understanding of the human microbiota landscape during transitions between states of health and disease.

We present eight HCT patients with three or more microbiologically-documented CDAD episodes, all of whom were successfully treated with FMT. For two of these patients, we were able to perform a longitudinal study of microbiota dynamics, employing shotgun metagenomic sequencing of serial stool samples obtained before, immediately after and over one year following FMT. We show that recipient stool assumed donor-like taxonomic and functional composition immediately following FMT, but that functional and taxonomic concordance were diminished after one year. These findings suggest that environmental, lifestyle and other host-factors are important determinants in long-term shaping of the microbiota. Furthermore, we show that despite the divergence of overall taxonomic and functional similarity at >1 year post-FMT, potentially important specific changes to the gene repertoire were retained, including the loss of pathogenicity genes from the pan-genome of *Escherichia coli* and a reduction of community-wide antibiotic-resistance genes.

## Results

### Treatment outcomes of FMT in eight HCT patients

FMT was delivered as an oral, encapsulated therapy using stool from a healthy unrelated donor (Table 1). All eight of the HCT patients to receive FMT experienced resolution of CDAD at 8 weeks, a standard time frame for determining cure. Subjects G and H died of underlying CNS disease and intracranial hemorrhage at 208 and 101 days following FMT, respectively. Both subjects were negative for CDAD at the time of death. Subjects A, B, C, E and F were free of CDAD at time of final stool sample collection at 408, 384, 456, 448 and 410 days post-FMT, respectively. Subject D experienced a recurrence of CDAD, and tested positive for CDAD at time of final stool sample collection, 179 days post-FMT. FMT is typically performed in patients with CDAD after significant clinical improvement is achieved with antimicrobial therapy. Subject B had active diarrhea at the time of FMT; the remaining five subjects were symptomatically quiescent. FMT was well tolerated in all subjects, and the subject with active diarrhea experienced symptom improvement within days of treatment. No attributable side effects or exacerbation of graft-versus-host disease were observed.

**Table I.**

| Subject | M/<br>F | Age at<br>FMT | Underlying<br>illness | Type of<br>HCT | CDAD history | Date of FMT<br>relative to HCT<br>(days post-HCT) | Rx | Symptoms at<br>time of FMT | Donor<br># | Recurrence of CDAD<br>post FMT at time of<br>latest sample<br>collection |
| --- | --- | --- | --- | --- | --- | --- | --- | --- | --- | --- |
| A | F | 56 | T-PLL | dUCB-HCT | 1,2,4 and 5 m<br>after HCT | 196 | V | BM character<br>declining off V;<br>no frank<br>diarrhea | 29 | No |
| B | F | 63 | DLBCL<br>with CNS<br>relapse | Auto-HCT | 1,2 m before and<br>1 m after HCT | 56 | V,Fi | 4-5 loose BM/d | 29 | No |
| C | F | 56 | Relapsed<br>AML | Allo-HCT (MRD) | Months after<br>second HCT with<br>8+ recurrences | 1160 | M,V,<br>Culturelle | 1 loose BM/d | 29 | No |
| D | F | 39 | AML | dUCB- HCT | 1 m before and<br>1,6 m after HCT | 577 | V | No significant<br>symptoms | 28 | No |
| E | M | 57 | CNS NHL | Auto--<br>HCT | 4 m after HCT,<br>and 6+<br>recurrences | 411 | V,<br>probiotics | No significant<br>symptoms | 29 | No |
| F | F | 38 | AML (FLT3<br>ITD+) | dUCB-HCT | 3 episodes starting<br>26 m after HCT | 926 | M,V | Frequent, formed<br>BM | 29 | No |
| G | M | 50 | DLBCL | Allo-HCT | 5 m before, 2 w<br>and 3 m after<br>HCT | 83 | V | 1-3 BM/d | 32 | N/A |
| H | M | 71 | ALL | Allo-HCT<br>(MUD) | 4 m after HCT | 129 | V | 3-4 loose BM/d | 34 | N/A |

### Short-term concordance and long-term divergence between recipient and donor microbiota following FMT

To better understand the taxonomic impact of FMT therapy in HCT recipients, we employed shotgun metagenomic sequencing of serial stool samples—both pre- and post-FMT—from two HCT recipients, and investigated bacterial genus, strain and gene-level changes over a period of 408 and 384 days for Subjects A and B, respectively. Sample collection for Subjects A and B was performed as follows: a single stool sample was obtained immediately before FMT; four stool samples were collected within a short-term period following FMT (on days 6, 14, 16 and 21 for Subject A; and days 6, 8, 13 and 20 for Subject B); and one stool sample was obtained at a long-term timepoint post FMT (days 408 and 384 for Subjects A and B, respectively). We also performed shotgun metagenomic sequencing of one long-term post-FMT sample from subjects C, D, E and F; pre-FMT samples were not available for these individuals. Long-term samples for subjects G and H were not collected as subjects were deceased at time of collection. Samples immediately pre-FMT and within 21 days post-FMT were not available for Subjects C, D, E, F, G and H.

In order to draw parallels between our results and the majority of FMT metagenomic studies to date, we began with an investigation of bacterial composition following FMT at the level of genus, a level of resolution typically targeted with the common approach of 16S rRNA gene amplicon analysis. For both Subjects A (Figure 1a) and B (Figure 1b) we observed a rapid shift towards a more donor-like state for all bacterial members within the most abundant genera. Although donor-concordant patterns of engraftment were found to be sustained for all short-term samples (<21 days), compositional similarity was largely diminished in the long-term (>1 year) timepoints for both Subjects A and B.

**Figure 1:**
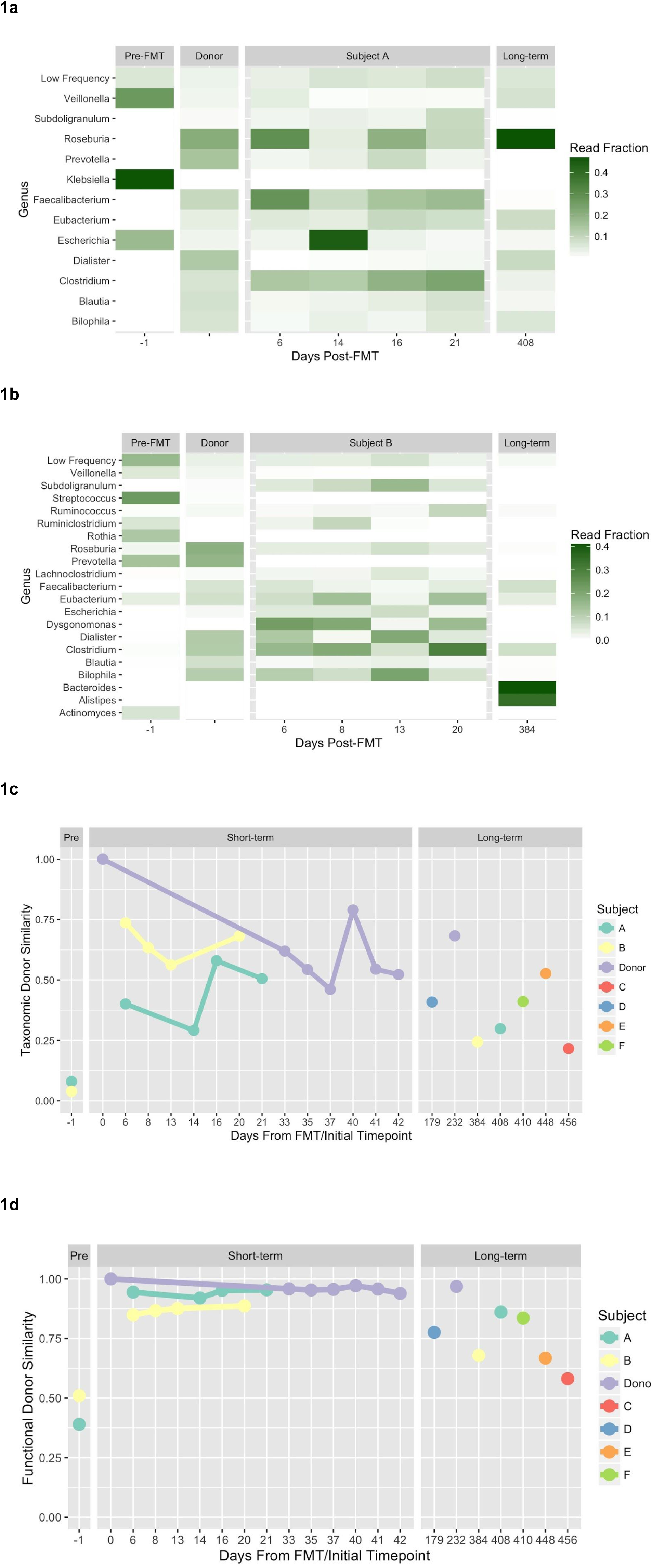
Genus-level taxonomic composition reveals extensive short-term donor microbial engraftment with FMT, followed by long-term reduction in donor similarity relative to donor self-similarity over a similar time period. Genera present in the donor appear in (a) Subject A and (b) Subject B recipients with inconsistent long-term residence. For visual clarity, genera represented by >5% of assigned reads from at least one timepoint are shown. (c) Whole-community Bray-Curtis donor similarity for Subjects A and B, and an additional four patients (Subjects C, D, E and F) for which long-term and donor samples were obtained, is shown for genus-level composition. Donor similarity is calculated for each recipient timepoint with the corresponding donor sample used for FMT. As a point of reference, we show multiple timepoints for Donor #29, where similarity is measured against the first timepoint. The variation in self-similarity within the short-term donor time series suggests an upper limit on potential recipient-donor similarity. Microbial sequencing reads were classified using Kraken (Wood and Salzberg 2014) in conjunction with a sequence database collected from NCBI Refseq and Genbank microbial genome references. (d) Whole-community donor Chao similarity in gene content shows similar patterns of donor similarity in the short-term and divergence in the long-term. Gene abundances were measured by alignment to the Uniref50 functionally annotated protein sequence database (Suzek et al. 2007).

To quantitatively assess parity between donor and recipient gut communities, we calculated whole-community similarity of all timepoints for Subjects A and B and long-term timepoints for Subjects C, D, E and F to corresponding donor timepoints at both the levels of genus (Figure 1c, Table S2) and gene (Figure 1d, Table S1). We observed a large increase in community-wide donor similarity at both the level of genus and gene in subjects A and B with FMT, with maintenance of elevated similarity during the three week short-term follow-up period. However, at >1 year post-FMT, donor similarity was reduced beyond the range of variability observed in donor self-similarity. As a point of comparison, we also calculated donor self-similarity using multiple stool samples taken over 180 days from donor #29, who provided fecal samples for all subjects aside from D, G and H. Donor self-similarity was calculated as similarity across the donor time series to the initial timepoint. At the taxonomic level, donor self-similarity fluctuated by as much as 33%, though at the level of gene, variations in donor self-similarity were minimal (within 1%). Although stool sampled immediately pre-FMT and within a month post-FMT were not collected for Subjects C, D, E, F, G or H, we were able to obtain samples for Subjects C, E and F at >1 year post-FMT, and for Subject D at ~6 months following FMT. We noted low donor similarity indices in these samples for both genus and gene (Figure 1c, d) (Table S1, S2).

### Strain-level genomic remodeling accompanies FMT-induced community-level turnover

To achieve a higher-resolution perspective of the metagenomic events surrounding FMT, we performed strain-level analysis on a subset of high-abundance taxa. Since the presence of genus Roseburia was found to be negligible in Subject A pre-FMT, yet abundant in both the donor and recipient post-FMT (Figure 1a), we sought to investigate the concordance of Roseburia strains between donor and Subject A over both the short-term (within 21 days of FMT) and the long-term (>1 year). Using nucleotide polymorphisms to distinguish between strains within the Roseburia genus, we observed the ratio of concordant to discordant single nucleotide variants (SNVs) to be similar between Subject A short-term samples post-FMT (Figure 2a), and those from the donor sampled from within a period of a month (Figure 2b). At >1 year post-FMT, we found the strains present within the genus Roseburia in Subject A to be genetically different from those of the donor.

**Figure 2:**
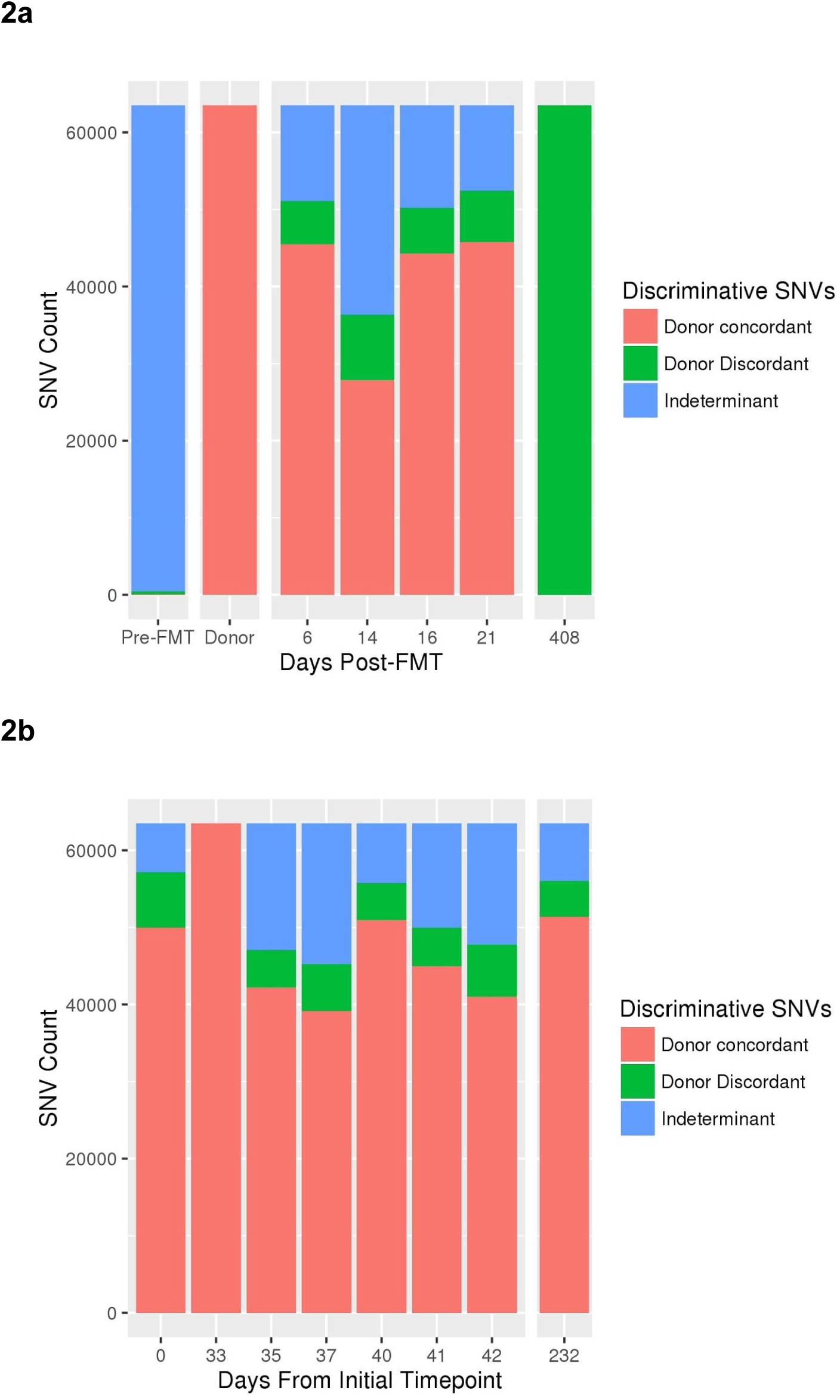
Dominant Roseburia strain(s) are different in the short versus long term timepoint after FMT. (**a**) A high number of single nucleotide variants (SNVs) distinguish the genus Roseburia strains present in the donor FMT sample (day 33 in donor time series) and Subject A recipient shortly after FMT from those in Subject A after one year following FMT. Roseburia was introduced with FMT, thus there were extremely few callable Roseburia SNVs in the pre-FMT timepoint. Thereafter, the relative proportion of donor-specific or long-term-specific SNV alleles tends heavily toward donor-specific during the short-term time points, indicating that the emergence of long-term discriminative nucleotide variation occurs in the period following the first three weeks post-FMT. (**b**) Roseburia strains demonstrate stable, low-level diversity in the donor across eight time points taken over ~8 months, contrasting with the divergence observed in the recipient. Sequences were assembled from short read data obtained at the long-term time-point, and resultant contigs were classified taxonomically (see methods). Contigs belonging to genus Roseburia were used as a reference for alignment of read data from all timepoints, followed by SNV calling and variable site selection. All polymorphisms distinguishing the donor from the long-term sample were selected, and presence of these discriminant polymorphisms in intermediate timepoints was recorded (adapted from a previously described approach (Li et al. 2016)).

As organisms from the genus Escherichia were observed in both donor and Subject A pre-FMT (Figure 1a), we focused on the representative species, *Escherichia coli*, and investigated whether strains post-FMT showed greater concordance with recipient pre-FMT or donor strains of *E. coli*. Using PanPhlAn (see Materials and Methods) to calculate individual gene abundances within the *E. coli* pan-genome, we observed large-scale changes in the gene complement of *E. coli* in the recipient following FMT (Figure 3). Hierarchical clustering identified 719 genes present prior to FMT that disappear within the first two timepoints post-FMT and remain absent through the >1 year timepoint (Figure 3, Supplementary Data File 1). This group is significantly enriched for known pathogenicity genes (Figure 3, Supplementary Data File 2). While there was a loss of many notable *E. coli*-specific virulence genes over time, there was an overall increase in the relative abundance of *E. coli* in the broader microbial community in Subject A post-FMT.

**Figure 3:**
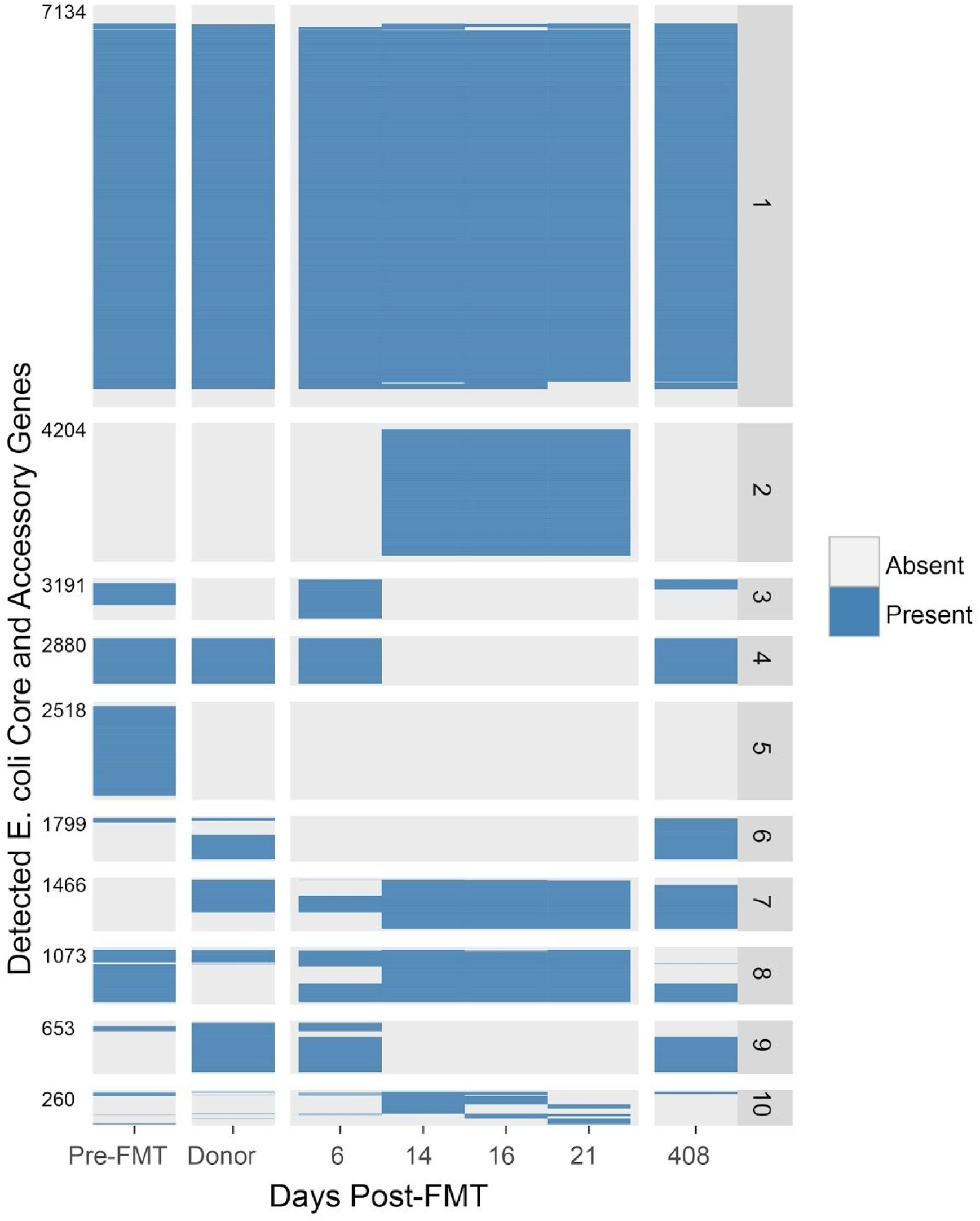
The total gene complement of *E. coli* in Subject A experiences large-scale remodeling in association with FMT, resulting in broad reductions in genes significantly enriched for virulence factors. Pronounced changes occur within the short-term timeframe, with further change in gene complement after one year. PanPhlAn (Scholz et al. 2016) was used to identify the presence of individual genes within the pan-genome of *E. coli*. Reads were aligned to a nonredundant collection of genes present in phylogenetically diverse *E. coli* isolate genome sequences. Those genes occurring at an abundance consistent with *E. coli*-specific occurrence were chosen and designated present or absent, as described previously (Scholz et al. 2016). Gene occurrence profiles were hierarchically clustered and divided into 10 groups with similar occurrence over the time series. Cumulative gene counts across groups are indicated on the left margin. Genes associated with virulence were identified by alignment to the Virulence Factor Database (Chen et al. 2005). Over- or underrepresentation of virulence genes in cluster groups was determined using Fisher’s exact test to detect significant departure from random occurrence. Significant (p < 0.05) underrepresentation was found in group 1, and significant overrepresentation was found in groups 5 and 9. KEGG annotations of genes within each group are given in supplementary data file 1. Virulence factors found within each group are given in supplementary data file 2.

To investigate the effects of FMT on the antibiotic resistome, we measured the abundance of antibiotic resistance genes from donor stool and Subject A both pre- and post-FMT (Figure 4). Subject B was excluded from this analysis due to a large concentration of human DNA pre-FMT, reducing the available microbial data volume. We performed alignment of shotgun data to the Comprehensive Antibiotic Resistance Database (McArthur et al. 2013) to obtain antibiotic resistance gene counts, normalized by total per-sample coverage and aggregated by resistance phenotype. Following FMT, we observed an immediate decrease in the abundance of antibiotic resistance genes by 2.4-fold which remained stable beyond 1 year. In addition, we noted a sustained increase in Chao similarity of unaggregated gene abundance profiles between the resistome profile of Subject A and the donor (Table S3).

**Figure 4:**
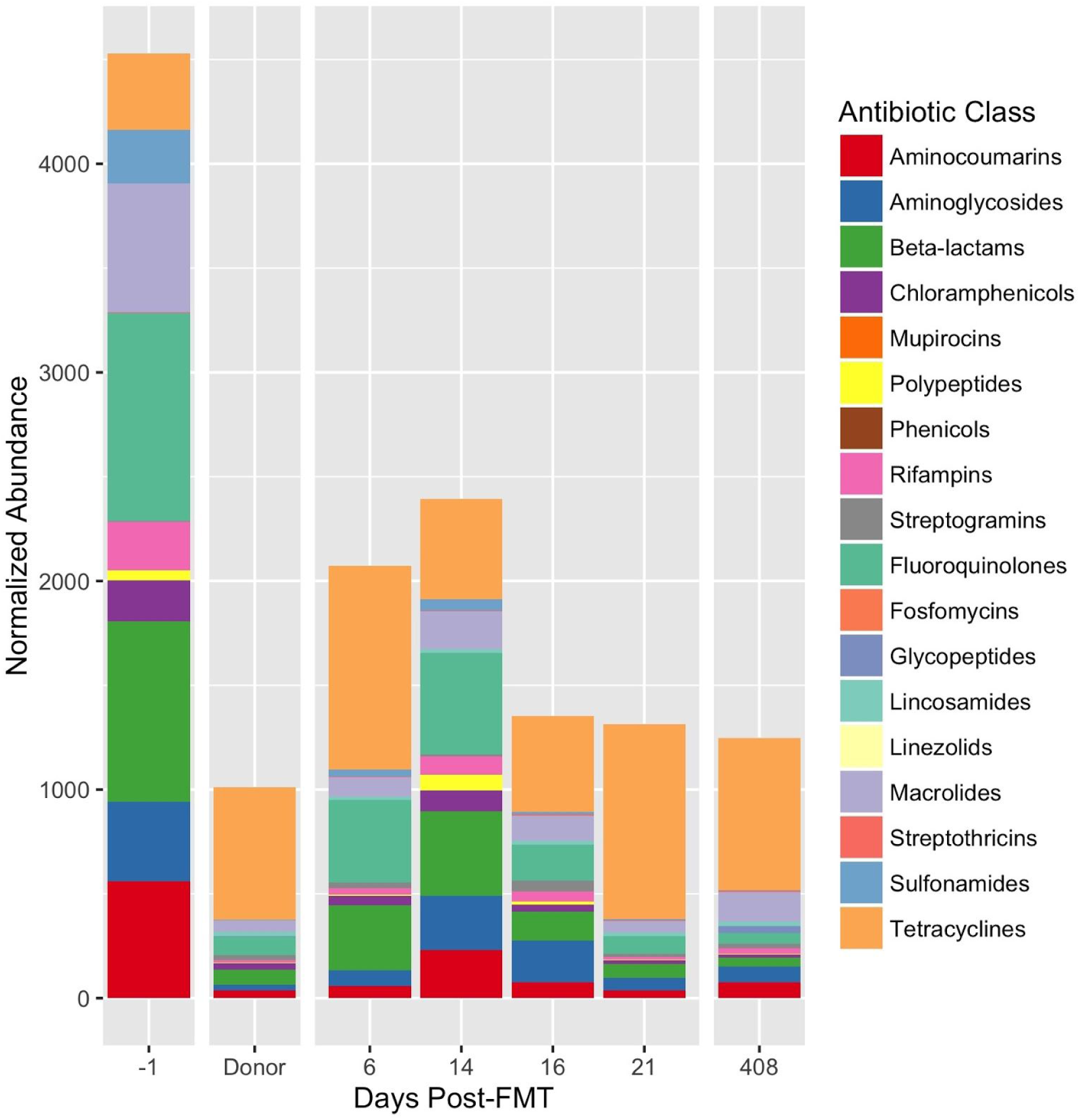
Antibiotic resistome profile of Subject A is rapidly remodeled immediately after FMT. Antibiotic resistance gene abundances were measured by alignment to the CARD antibiotic resistance database and normalizing by per-sample coverage. Individual gene abundances are aggregated by antibiotic class. Genes conferring multiple antibiotic class resistance phenotypes are counted toward each antibiotic class.

## Discussion

Our study provides one of the first metagenomic analysis of strain- and gene-level bacterial engraftment following FMT, and the first such analysis in patients being treated for CDAD. The success of FMT in resolving CDAD in HCT patients has been demonstrated previously (Kelly et al. 2014; Mittal et al. 2015; Neemann et al. 2012; Aroniadis et al. 2016; Webb et al. 2016), however a detailed characterization of this “black box” therapy has been lacking. We used a shotgun metagenomic approach to examine both short- and long-term bacterial engraftment following FMT to treat CDAD in HCT patients. We show that although concordance was observed between microbiota composition of the donor and recipient in the short-term, concordance was reduced after 1 year. Previous studies investigating long-term donor similarity have been confined to immunocompetent individuals and have either (1) been limited to 16S rRNA sequencing analysis (Broecker et al. 2016; Jalanka et al. 2016) or (2) observed strain-level changes, but only through to three months following FMT. (Li et al. 2016)

In the case of the 16S rRNA studies, Broecker et al. (Broecker et al. 2016) investigated family donor FMT in one subject, and reported long-term (>4.5 years) donor similarity but limited short-term (<7 months) concordance. Conversely, we observed short-term similarity and long-term donor discordance. For all eight HCT patients in our study, recipients received stool from a healthy, unrelated donor, in contrast to the method employed by Broecker et al, where fecal material was donated by the sister of the FMT recipient. In light of our findings that unrelated donors and recipients display dissimilar microbiota profiles at long-term timepoints post-FMT, we postulate that FMT engraftment is broadly impacted by external effects such as diet, host genetics, host immune surveillance, and subsequent antibiotic exposure, which may supersede FMT composition in determining long-term FMT durability. If true, then the Broecker study may simply demonstrate microbiota convergence towards a donor-like state when external factors are common between recipient and donor, complementing our observation of divergence when they presumably differ. Lifestyle and diet questionnaires are not currently standard practice in long-term metagenomic sample collection, and both our study and that of Broecker et al. lack the data to test the effects of external factors such as antibiotic exposure, lifestyle and diet on gut microbiota composition after FMT. Should such a nuanced and multifaceted perspective of engraftment prove accurate, then attempts to alter the microbiota in a predictable and sustained fashion will require knowledge and potentially manipulation of factors beyond the microbial composition of FMT.

Also employing 16S rRNA sequencing, Jalanka et al. used FMT from “universal donors” to treat CDAD in 14 immunocompetent patients, and report on bacterial composition both in the short-term (3 days, 2 weeks and 1 month) through to 1 year following FMT (Jalanka et al. 2016). Like Broecker et al., Jalanka et al. found the unrelated donor bacterial profile to be retained in recipients at 1 year post-FMT. These observations of long-term donor similarity in immunocompetent individuals contrast with our findings of long-term donor dissimilarity in HCT patients. We attribute this contrast to differences in sensitivity between 16S and metagenomic methodologies, and to possible involvement of the host immune status and potential subsequent therapies in influencing the long-term fate of donor microbiota.

Our analysis of nucleotide-level similarity between donor and recipient, adapted from Li et al (Li et al. 2016), showed donor concordance over short-term timepoints consistent with that group’s findings, however after 1 year post-FMT we report high discordance between donor and recipient genus Roseburia SNVs (Figure 2). We focused on this genus since it appeared to be absent in the recipient pre-FMT, abundant in both the donor and the recipient at all timepoints post-FMT, and showed the highest abundance of reads at >1 year post-FMT. Given the consistent presence of Roseburia across all timepoints, it was surprising that the genomic sequence(s) of Roseburia at >1 year post-FMT was extensively dissimilar to that of the Roseburia organism(s) that had initially engrafted in the recipient. This suggests that ecological niche availability within the gut environment may inform taxonomic composition, but intra-species competition, neutral drift, or other mechanisms of nucleotide divergence continue to drive strain turnover within a given niche. Indeed, the theory of ecological niche availability within the gut driving microbiota composition is not new (Costello et al. 2012). More studies are needed to understand the importance of sub-species interactions and how such fluctuations might affect gut microbiome behavior and ultimately human health.

To gain a better appreciation for how gene-level differences between same-species strains of bacteria affect engraftment following FMT, we chose to investigate the gene complement of *E. coli*, a species that exhibits wide variability between strains, ranging from dominant commensal in the human gut to highly virulent pathogen (Tenaillon et al. 2010). We chose this organism both because it occurs at high abundance through the time series obtained from Subject A, and because there is an extensive genome collection available for *E. coli* with which to study the accessory gene pool of this organism. At the taxonomic level, the presence of *E. coli* was consistent in Subject A throughout all samples; however at the gene level we observed massive losses and gains in the total *E. coli* accessory genome between all measured timepoints, with the most pronounced differences occurring pre- and post-FMT. Whether this change is due to strain replacement or genomic remodeling within a single resident strain is unconfirmed, although the rapid timeframe in which it occurs suggests the former.

The group of *E. coli* genes present before FMT and lost shortly following FMT were found to be enriched in virulence genes (Figure 3). These genes encompass a range of virulence mechanisms such as siderophore biosynthesis, enterotoxin production, type III secretion systems, and adhesin proteins (for a complete list of genes see supplementary data files 1 and 2). The disappearance of genes following FMT suggests that, in addition to being a viable treatment for CDAD, FMT may also hold potential as a treatment for certain infections caused by virulent *E. coli*; for instance, disease caused by Shiga toxin-producing *E. coli*, which is known to be exacerbated by certain antibiotics and is therefore difficult to treat conventionally (McGannon et al. 2010). The case of FMT-mediated loss of virulence genes from *E. coli* is an example of how a shotgun metagenomic approach to studying FMT can enhance not only our understanding of strain-level dynamics post-FMT, but also supports other treatment applications beyond *C. difficile* infection.

Multi-drug-resistant (MDR) bacteria are becoming increasingly common in patients with hematological malignancies, such as HCT recipients (Baker and Satlin 2016). It has been previously shown that FMT is capable of reducing antibiotic resistance genes in immunocompetent patients being treated for CDAD (Millan et al. 2016). Due presumably to both the requirement that antibiotic treatment be administered for at least 3 CDAD episodes before FMT will be attempted, as well as the use of antibiotics in conjunction with HCT therapy, recipients have been found to have higher numbers of antibiotic resistance genes than those of healthy donors. We sought to determine (1) whether immunocompromised patients carry an increased number of antibiotic resistance genes compared to a healthy donor, and (2) if FMT is similarly capable of reducing the abundance and variety of antibiotic resistance genes in HCT patients. We demonstrate that, for the immunocompromised subject time series analyzed, there was an increased presence of antibiotic resistance genes in the recipient compared to the donor, and that abundance of the antibiotic resistome was reduced by >50% followed FMT (Figure 4). Furthermore, this reduction persisted more than one year after FMT. This is consistent with findings in immunocompetent recipients, and bodes well for the potential reduction of antibiotic resistance via FMT in HCT patients. Given the lean toolkit of antibiotics that remain effective in the face of MDR pathogens, identifying strategies to mitigate antibiotic resistance is of tremendous importance, especially for immunocompromised individuals. Reducing antibiotic resistance may prove to be yet another potential application of FMT.

In conclusion, our work establishes a foundation for future studies employing a shotgun metagenomic approach to better understand the evolution of microbial communities after FMT in various disease states. We demonstrate that granular shotgun metagenomic analyses show high initial donor similarity and long-term donor divergence at the community-wide taxonomic and functional levels, as well as in single-species gene composition and nucleotide variation. These results lend support to the safe use of FMT for CDAD and reveal short and long-term FMT dynamics in immunocompromised HCT patients. In addition, these results strongly support the value of detailed analysis of metagenomic sequence data, and suggest a holistic model of FMT engraftment durability informed by more than donor or recipient microbiome taxonomic composition at the species, genus or phylum level.

## Methods

### Stool sample collection

BMT-recipient fecal samples were collected by patients at home using collection kits provided. Fecal samples were shipped to the laboratory on pre-frozen freezer packs and stored at −80°C within 24 hours of production. Upon defecation, pre-FMT and early (<21 days) timepoint fecal samples were aliquoted in 95% EtOH. Donor and all late timepoint (>6 months) fecal samples were stored in the absence of EtOH.

### Fecal microbiota transplantation

Donors were screened and fecal samples were collected, and processed into FMT capsules (Capsugel hypromellose capsules) as described (Youngster et al. 2014, 2016; Hamilton et al. 2012). FMT was performed as previously reported (Youngster et al. 2014). Briefly, each patient received 15 frozen FMT capsules with water on each of two consecutive days followed by monitoring for any adverse effects. Subjects took nothing by mouth for four hours before and one hour after administration. Subjects were instructed to stop oral vancomycin 24-48 hours prior to FMT. Patient A vomited more than 12 hours after the first dose but did not bring up any capsules. There were no serious adverse events attributable to FMT.

### DNA extraction

Stool samples were stored at −80°C until use. DNA extractions were performed on all samples using the QIAamp DNA Stool Mini Kit (QIAGEN^®^) as per manufacturer instructions, with an added bead-beating step using the Mini-Beadbeater-16 (BioSpec Products) and 1 mm diameter Zirconia/Silica beads (BioSpec Products) consisting of 7 rounds of alternating 30 second bead-beating bursts followed by cooling on ice. DNA concentration and quality estimations were performed using Qubit^®^ Fluorometric Quantitation (Life Technologies) and Bioanalyzer 2100 (Agilent). DNA libraries were prepared using the Nextera XT DNA Library Prep Kit (Illumina^®^). Prepared libraries were subjected to 100 base pair, paired-end sequencing on a HiSeq 2500 (Illumina^®^).

### Metagenomic Shotgun Sequencing and Analysis

Data from individual sequencing runs were first aggregated by clinical sample using an in-house script. Paired-end raw reads from shotgun sequencing were trimmed using cutAdapt v1.7.1 (Martin 2011) using a minimum length 80bp, maximum N count of 0, minimum terminal base score of 30, and the Illumina Nextera transposase and adapter sequences. In addition, reads were trimmed by 16bp at the 5’ end to remove low quality bases and deduplicated with PrinSeq v0.20.4 (Schmieder and Edwards 2011). Sequencing was performed to asymptotic saturation of available taxonomic diversity, assessed by calculating rarefaction curves using the vegan v2.4 (Dixon 2003)package in the R statistical computing language (Ihaka and Gentleman 1996) and defining an asymptotic threshold at less than one new genus discovered per each additional million reads (Figure S1).

Reads were taxonomically assigned using Kraken v0.10.4 (Wood and Salzberg 2014) based on k-mer resemblance to human, bacterial, viral and fungal k-mer profiles (k = 31) generated from the NCBI Bacterial Refseq and Genbank collections (accessed November 14, 2015), filtered with the kraken-filter utility at a threshold of 0.2, and results visualized with in-house scripts. Similarity between samples based on taxonomic calls was calculated with the Bray-Curtis similarity metric (Beals 1984).

Functional analysis was conducted on quality filtered reads using HUMAnN2 v0.2.1 (Abubucker et al. 2012) to obtain read alignments to the Uniref50 functionally annotated protein sequence database (Suzek et al. 2007). Relative gene family abundances were obtained by dividing gene family coverage by per-sample total coverage. The Chao similarity metric was chosen to assess functional similarity due to the high diversity present in the functional repertoire (~2 million distinct genes) and the consequent risk of negative bias due to undersampling, a problem affecting alternative choices of similarity index such as Bray-Curtis or Jaccard. (Chao et al. 2005)

Roseburia SNV diversity was calculated by first assembling reads from the subject 1 long-term time-point with Spades v3.6.1 (Bankevich et al. 2012-5), then aligning reads from all timepoints to the contigs so obtained using BWA (Li and Durbin 2009), and finally calling SNVs using the GATK Unified Genotyper (Van der Auwera et al. 2013). SNVs discriminating donor from long-term recipient timepoint were selected as previously described (Li et al. 2016) and visualized with an in-house script.

PanPhlAn (Scholz et al. 2016) was used to obtain gene repertoire membership information for *Escherichia coli* using the 2016 pan-genome index for this organism. Gene functions were annotated for all genes using the KEGG (Kanehisa and Goto 2000) database according to the method described in the PanPhlAn documentation. Those genes occurring at an abundance consistent with *E. coli*-specific occurrence were chosen and designated present or absent as described previously (Scholz et al. 2016). Genes associated with virulence were identified by alignment to the Virulence Factor Database (Chen et al. 2005). Over- or underrepresentation of virulence genes in cluster groups was determined using Fisher’s exact test to detect significant departure from random occurrence.

Antibiotic resistance profiles were obtained by using BWA (Li and Durbin 2009) to align read data from all samples to the Comprehensive Antibiotic Resistance Database (McArthur et al. 2013) (accessed 6/6/16) and collecting per-gene read counts and aggregating by resistance phenotypes using an in-house script.

Plots for all analyses were generated in the R statistical language (Ihaka and Gentleman 1996) using the ggplot2 (Wickham 2009), reshape2 (Wickham 2012), and doBy (Højsgaard et al. 2006) packages.

## Additional Information

### Data Deposition and Access

The sequence data from this publication have been submitted to the NCBI SRA database under BioProject accession No. PRJNA349197. All code written for the analysis and visualization of this work are included in Supplementary Data File 4.

### Ethics Statement

Patients were identified and referred for FMT by hematologists or oncologists, and the study was reviewed and approved by the Partners Human Research Committee (Boston, MA); subjects provided written informed consent. The project was supervised by the FDA under IND 16011 held by E. Hohmann.

## Author contributions

ASB and ELH conceived of the study; ELM, SBF and ASB wrote the manuscript; SBF and ET performed the molecular biological assays described in the manuscript; ELM and MW performed informatic analyses; HS and JM managed patient correspondence, sample collection and sample handling; ELH designed and carried out the clinical intervention described in this manuscript; DAR provided detailed feedback and commentary on the interpretation of results and the manuscript.

## List of abbreviations

FMT (fecal microbiota transplantation); CDAD (*Clostridium difficile*-associated disease); F (female); M (male); T-PLL (T prolymphocytic leukemia); DLBCL (diffuse large B-cell lymphoma); CNS (central nervous system); AML (acute myelogenous leukemia); NHL (non-Hodgkin lymphoma); FLT3 ITD (FLT3 internal tandem duplication); ALL (acute lymphoblastic leukemia); dUCB (double umbilical cord blood); HCT (stem cell transplantation); Auto (autogenic); Allo (allogenic); V (vancomycin); Fi (fidaxomycin); M (metronidazole); BM (bowel movement); d (day); w (week). N/A is indicated for patients who were deceased prior to the time of final long term timepoint stool sample collection.

